# Composite interval mapping and genomic prediction of nut quality traits in American and American-European interspecific hybrid hazelnutss

**DOI:** 10.1101/2025.10.20.683460

**Authors:** Scott H. Brainard, Julie C. Dawson

## Abstract

The native, perennial shrub American hazelnut (*Corylus americana*) is cultivated in the Midwestern U.S. for its significant ecological benefits, as well as its high-value nut crop. Genetic improvement of perennial crops involves long-term breeding efforts, and benefits from the use of genetic data in selection to reduce breeding cycle time. In addition, high-throughput phenotyping methods are essential to the efficient and accurate screening of large breeding populations. This study reports novel advances in both of these domains, for American (*C. americana*) and interspecific hybrids between European (*C. avellana*) and American hazelnuts. Two populations of hazelnuts, one composed of *C. americana* and one composed of *C. americana* x *C. avellana* hybrids, were phenotyped over the course of two years in two locations using a digital imagery-based method for quantifying morphological nut and kernel traits. This data was used to perform composite interval mapping (CIM) using a recently released genetic map, and genomic prediction using a newly-available chromosome-scale reference genome for *C. americana*. Multiple QTL were detected for all traits analyzed, with an average total R^2^ of 52%. Genomic prediction exhibited high accuracy, with an average correlation coefficient between genotypic values and phenotypic observations of 0.78 across both environments. These results suggest that incorporating genetic data in selection is a tenable method for improving genetic gain for highly-polygenic traits in hazelnut breeding programs.

**Core ideas:**

- Morphological nut characteristics are under polygenic control in American and American- European interspecific hazelnuts.
- Best linear unbiased predictors allow for accurate prediction of morphological nut characteristics.
- Marker density and training population design must be tailored to the sample population for which predictions are being made.

## INTRODUCTION

Hazelnuts (*Corylus* spp.) are a globally significant nut crop, with annual production of over 1.3 million tons, produced across 34 countries (FAOSTAT, 2023). Currently, this cultivation is limited to areas with climatic conditions approximating the Mediterranean region in which the dominant cultivars of *C. avellana* were bred (Mehlenbacher and Molnar, 2021). The narrow range of cultivars has limited U.S. production of hazelnuts to the Willamette Valley in Oregon, where over 95% of U.S. acreage is situated (USDA Economic Research Service, 2025). The native species *C. americana*, which is endemic and well-adapted to much of the eastern U.S., represents a valuable source of genetic diversity for expanding hazelnut production. American hazelnut can exhibit vigorous growth in cold environments, and robust resistance to endemic pathogens such as Eastern Filbert Blight (EFB, *Anisogramma anomala*) (Molnar et al., 2018; Revord et al., 2020).

In this context, the Upper Midwest Hazelnut Development Initiative (UMHDI) was established to develop regionally adapted hazelnut cultivars for the Upper Midwest. The program’s long- term goal is to establish a sustainable, cold-hardy, disease-resistant Midwestern hazelnut industry that complements that of Oregon. However, the genetic improvement of hazelnut is limited by its long generation time. While the most economically important traits for growers are total kernel yield, kernel size, and kernel percentage, as these directly determine processing efficiency and market value, a seedling hazelnut will not typically flower under field conditions for at least three years. In addition, mature plant yields will often not be fully apparent for a decade, and multi-year harvest data are necessary to precisely quantify such traits. Methods that can make parental and progeny selection decisions more efficient are therefore extremely valuable for a perennial species such as hazelnut, by accelerating the traditional breeding process.

To overcome the extended generation time and delayed trait expression inherent to perennial species, this study evaluates several complementary approaches—linkage mapping, genomic prediction, and high-throughput phenotyping—to enable earlier, more precise selection in hazelnut breeding programs. Quantitative genetic tools are essential in this regard. The identification of QTL is essential to the implementation of marker-assisted selection, while genomic prediction can dramatically accelerate genetic gain for polygenic traits (Würschum, 2012; Heslot et al., 2015). The recent development of genomic resources for *C. americana* have made both of these approaches accessible to breeding programs. First, a genetic map constructed using an F_1_ progeny family descended from a cross between Oregon and Midwestern hazelnut varieties ‘Jefferson’ and ‘Eric4-21’, respectively, now permits linkage map-based approaches to detecting QTL (Brainard et al., 2023). Such interspecific populations are highly relevant to Midwest breeding efforts, as *C. avellana* remains a valuable source of diversity for nut quality-related traits. The use of linkage mapping populations is well-suited to such populations, where the contrasting phenotypes of parental varieties frequently produces high degrees of segregation in progeny families. In addition, chromosome-scale genome assemblies for the *C. americana* selections ‘Rush’ and ‘Winkler’ also now allow for the efficient identification of polymorphisms for use in genomic selection models (Brainard et al., 2024).

This study uses both of these tools, in combination with a novel digital imagery-based phenotyping pipeline for precisely quantifying morphological characteristics of in-shell hazelnuts and kernels. Recent developments in contour analysis of digital images has led to numerous methods for quantifying the sizes and shapes of plant structures (Hameed et al., 2018). Morphological features of in-shell nuts and kernels certainly do not comprise a comprehensive set of phenotypes that are important in hazelnut breeding. However, by focusing on a limited set of traits with high heritability, that are amenable to digital image–based phenotyping, we were able to make precise comparisons between QTL mapping and genomic prediction methodologies. To do so, we utilized two experimental populations. First, a wild population of *C. americana* sourced from the Wisconsin Department of Natural Resources was used as a “diversity panel”. Diversity panels are useful for assessing highly polygenic traits, and developing such a resource for American hazelnut is appropriate, as this native species not only has potential as a crop for the Upper Midwest, but is also a useful source of traits such as resistance to Eastern Filbert Blight, and cold hardiness. Second, F_1_ interspecific biparental progeny families generated by the University of Minnesota hazelnut breeding program were used to perform linkage mapping. The first population allowed for the evaluation of genomic prediction models to estimate trait heritability and assess the potential for early selection in hazelnut breeding, while the second population facilitated the identification of QTL associated with kernel traits in a *C. americana × C. avellana* F_1_ family.

## MATERIALS AND METHODS

### Plant Materials

Two populations of seedling hazelnut plants were used in this study. One was composed of 473 wild *C. americana* seedlings sourced from the Wisconsin Department of Natural Resources and planted in 2016 at a farm outside of Barneveld, Wisconsin (43.043266 N, −89.926068 W). This population was treated as a “diversity panel”, and used to fit and validate genomic prediction models that could be used when introgressing valuable genetic diversity from *C. americana* into Midwest breeding programs. The second was composed of 258 plants which belong to three F_1_ families descended from crosses between clonal varieties sourced from the University of Minnesota and Oregon State University, specifically: Eric4-21 x Jefferson (“Eric\Jeff”), Gibs5- 15 x OSU-919-031 (“Gibs\OSU”), and Gibs5-15 x York (“Gibs\York”). These were planted at a University of Minnesota research farm in Rosemount, MN (44.695784 N, −93.079196 W) in 2015 and 2016. These represented the interspecific biparental families that were used to test both genomic prediction in breeding populations that include *C. avellana*, as well as perform F_1_ linkage mapping.

### Phenotyping

Morphological characteristics of in-shell hazelnuts and shelled kernels were used as the trait data for this study. This was collected primarily using an adapted version of the digital imagery acquisition and analysis pipeline reported by (Brainard et al., 2021) (Fig. 1). In brief, bushes were completely harvested by hand in August 2020 and August 2021. Harvested clusters were dried in the greenhouse and husked. A sample of 30 in-shell nuts were randomly sampled per bush, and weighed in bulk. These nuts were arranged on a 6x5 grid with a QR code, and a Nikon 5600 DSLR camera tethered to a desktop computer was used to acquire a single image. The OpenCV Python library was used to isolate each in-shell nut, and produce a binary mask by applying a fixed hue-saturation-value threshold to each pixel. An ellipse was then fit to each binary mask, and the length of its major (termed “length”) and minor axis (termed “width”) was calculated by converting pixel length to physical distance using a scale bar embedded in each image. Circularity of the nut was calculated as the ratio of these two lengths. The “height” of each nut along the axis perpendicular to the 2D photo was then measured using digital calipers; this height measurement is alternatively called “depth” in other studies (Yao and Mehlenbacher, 2000). Each nut was individually cracked, and the kernel was then returned to the grid, preserving the original arrangement of the nuts. A second photo was acquired, and the same traits were calculated, including caliper measurements of individual kernel heights. Finally, the kernels were weighed in bulk. This allowed for both a volumetric and gravimetric estimation of the percent kernel for each nut sampled. Python scripts for the image acquisition, processing, phenotyping, and file management are available at: https://github.com/shbrainard/hazelnut-phenotyping. Prior to large-scale deployment, the image-based phenotyping pipeline was validated against hand-measured reference data for in-shell and kernel length and width from 200 individual nuts, which confirmed extremely close correlations between manual and digital measurements (Supp. Fig. 1). With a RMSE < 0.4 mm for all four traits, the digital platform was judged to be extremely accurate, as confirmed previously by Brainard et al. (2021). Phenotypic measurements were averaged for each individual plant. The prcomp R function was used to perform PCA on this phenotypic data, and the ggplot2 function autoplot was used to visualize the relationship between these traits. All phenotypic data acquired via this pipeline is available via DataDryad: https://doi.org/10.5061/dryad.ghx3ffc0z.

**Figure 1.**
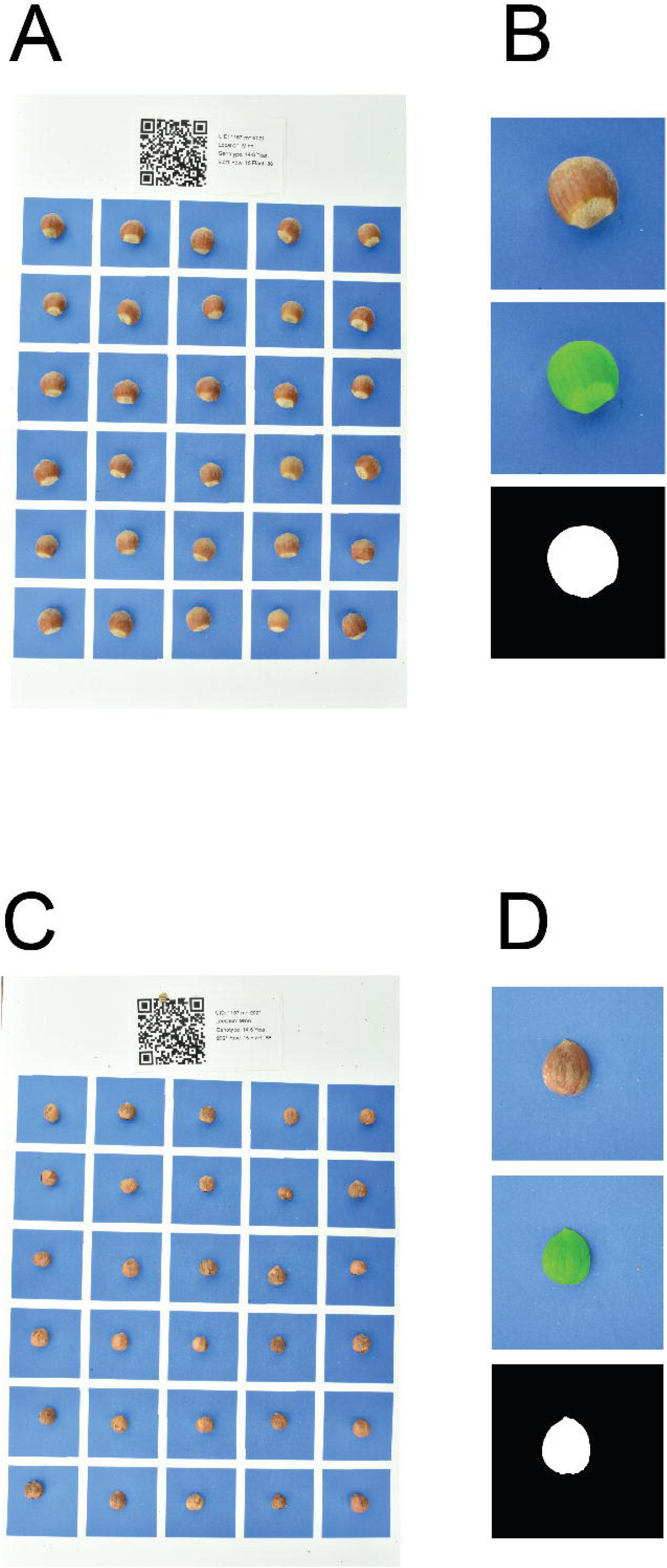
Digital image acquisition and processing pipeline. Staged images of subsamples of 30 in-shell nuts (**A**) and kernels (**B**) were acquired using a tethered DSLR camera. QR codes were scanned, with identifying information used for file management. Blue boxes containing individual nuts or kernels were then cropped, with the dimensions of each box being used as a scale for converting pixel resolution to spatial resolution. HSV indexing was used to create a binary mask of each nut or kernel, with bounding boxes fit to each mask to obtain phenotypic measures.

### Sequencing

Roughly 1 cm^2^ of leaf tissue was sampled from each bush in May 2021, immediately following budbreak. Tissue was sampled into 96-well Qiagen Collection Microtubes (Qiagen N.V., Venlo, The Netherlands), and lyophilized using a Labconco 18 L freeze dryer set to 0.004mBar for 72 hours. Freeze-dried tissue was then macerated. DNA extraction, quantification, library preparation, and sequencing was performed at the University of Wisconsin-Madison Biotechnology Center. Libraries were prepared for genotyping-by-sequencing using a double digestion with the restriction enzymes *NsiI* and *BfaI* following the methodology described by Elshire et al., 2011. This combination was pre-selected based on an analysis of k-mer distributions of various enzyme digestions, wherein *NsiI/BfaI* was observed to maximize the k-mer diversity of the library. Illumina adapters and sample-specific barcodes were then annealed. Samples were multiplexed, and paired-end 150-bp sequence data was generated using an Illumina NovaSeq 6000, with an average of 10 million reads per sample. The use of GBS was selected to mitigate the potential for ascertainment bias, compared with fixed-site genotyping platforms such as SNP arrays or amplicon sequencing, because loci are discovered *de novo* in each dataset rather than preselected from a reference population. Trimming and demultiplexing of raw Illumina reads was performed using a custom Java application https://github.com/shbrainard/gbsTools. Reads were aligned to the *C. americana* genome for ‘Winkler’ (Brainard et al., 2024). This genome was selected to again minimize ascertainment bias, as this assembly has previously been found to be structurally similar to published *C. avellana* reference genomes (Brainard et al., 2024). Due to its high quality, Winkler yielded the greatest proportion of high-quality read alignments of all tested assemblies, indicating broad suitability for aligning reads from both *C. americana* and interspecific hybrid plant material.

### SNP calling

Haplotype-based markers were called using Stacks 2 (Rochette et al., 2019), which can identify multiallelic markers using the phased nature of multiple indels or SNPs that appear within a single 150-bp paired-end read. Because such markers cannot be directly filtered for depth, the parameter ‘gt-alpha’ was increased to 0.01 as a method for ensuring genotype quality. Increasing this value allows for an alternative to filtering on depth in the resulting VCF, increasing confidence in accurate genotype calls for multiallelic loci. Markers were then filtered for linkage disequilibrium using bcftools (using the +prune plugin with a window size of 10kb, and r^2^ filter of 0.95, retaining only one site per window with the highest minor allele frequency (MAF)). This generated a set of 78,079 markers, with an average of ∼7,000 per chromosome.

For the purpose of genomic prediction, biallelic SNPs were called using the TASSEL GBSv2 pipeline (Bradbury et al., 2007). We considered the use of multiallelic markers for this analysis as well, but comparison of genomic relationship matrices derived from the Stacks (as described above) and TASSEL (as described below) pipelines showed a high degree of concordance (r² = 0.96), indicating that both marker types captured nearly identical patterns of relatedness within the populations analyzed. The 3,792,202 SNPs which were called were first filtered to exclude sites where the 80^th^ percentile of allele depth across all samples was > 8, leaving 1,112,817 sites. SNPs were then filtered to remove sites where minor allele frequency was < 0.05, or sites with more than one alternate allele, resulting in 80,852 sites. Samples were then filtered to remove individuals with greater than 5% missingness across all sites. Next, filtering was performed for linkage disequilibrium (LD) (r^2^ < 0.75, with the same selection criteria as above), which retained 52,402 SNPs. These thresholds were selected to balance high marker quality, with sufficient genome-wide coverage for estimation of relatedness.

Finally, markers were then subset to only include those which were retained in both the *C. americana* diversity panel in Wisconsin and interspecific biparental populations in Minnesota, leaving 50,961 SNPs. All filtering was performed using bcftools (Danecek et al., 2021). Because the impact of MAF and LD filtering can be affected by population structure, we compared several alternative filtering strategies in which variant filtering for depth, MAF, and LD was performed either on the full dataset, or separately within each population. Although the total number of retained SNPs varied among these approaches, subsequent genomic prediction analyses (as described below) showed negligible differences in ultimate prediction accuracy.

### Linkage map construction and composite interval mapping

Since the interspecific biparental populations in Minnesota were constructed from controlled crosses between known hazelnut varieties, it was possible to build a genetic map by using the R package onemap (Margarido et al., 2007) (https://github.com/augusto-garcia/onemap). This map was previously reported in (Brainard et al., 2023). Briefly, markers called using Stacks 2 were first filtered to include only those of segregation types A1, A2 and B3.7 (following the notation of Wu et al. (2002)), such that only markers with either three or four alleles remained. In these segregation types, the four possible alleles present in a biparental cross between two outbred diploid parents are represented as *a, b, c,* and *d*. A1 segregation types are loci for which the parental genotypes are *ab* x *cd*, A2 represents *ab* x *ac* crosses, and B3.7 represents *ab* x *ab*. This approach can be distinguished from previous maps built in F_1_ hazelnut populations, which have used only biallelic markers, or treated multiallelic markers as dominant markers (Torello Marinoni et al., 2018). Next, markers for which more than 5% of all samples had no called genotype were removed and two-point recombination frequencies were calculated for all possible phase configurations between all remaining markers using maximum likelihood. Maximum likelihood estimates were able to fully resolve phase between pairs of markers, due to the fully-informative segregation types that were used. A hierarchical clustering algorithm was used to construct linkage groups, and markers were ordered and phased within these groups by using a Hidden Markov Model with an error rate of 0.05. Recombination frequencies were then converted to genetic distances using the Kosambi mapping function (Kosambi, 1943). Finally, this map was imported into the R package fullsibQTL (Gazaffi et al., 2020) (https://github.com/augusto-garcia/fullsibQTL) which was used to perform composite interval mapping. Cofactors were selected to minimize the Akaike Information Criteria, and composite interval mapping was employed to perform a single QTL scan for each of the phenotypes measured using the digital image analysis protocol described above. Permutation tests were calculated to empirically estimate a LOD threshold representing a 0.05 genome-wide *p*-value. Additive and dominance effects were estimated for QTL exceeding this threshold using the function cim_char(). Least squares estimation was used to calculate the percent variance explained for both these QTL individually, and collectively, using the function *r2_ls*. This *R^2^* was calculated using the residual sums of squares (RSS) for the null and full models:

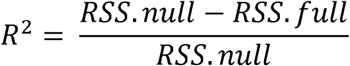

### Calculation of GEBVs

In order to calculate genomic-estimated breeding values from the biallelic SNP dataset described above, the R package StageWise (Endelman, 2023) (https://github.com/jendelman/StageWise) was used to compute variance components and best linear unbiased predictors (BLUPs) of additive genetic value. This software is designed to perform two-stage analysis, by first computing best linear unbiased estimators (BLUEs) for each genotype using a specified experimental design. Since the genotypes in both populations were comprised of unreplicated seedlings, a fixed-effects linear model was used to first compute BLUEs for each genotype where:

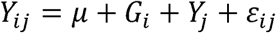

𝐺_𝑖_ represents the *i*^th^ genotype effect, 𝑌_𝑗_ the *j*^th^ year effect, and 𝜀_𝑖𝑗_ the residual variance (which included the 𝐺 ∗ 𝑌 effect), with 𝜀_𝑖𝑗_ ∼ 𝑁(0, 𝜎 ^2^)). Models were fit for each trait independently, and then passed to the function Stage2 in order to estimate genotypic values for each trait individually. Additive genetic values are here assumed to follow a multivariate normal distribution with a variance-covariance matrix proportional to the G matrix (VanRaden, 2008), where a centered marker of allele dosages **W** is normalized by the product of the major and minor allele frequencies (*p* and *q*) for the *k*^th^ marker:

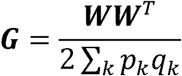

This model was selected after comparing the percent variance explained (PVE) of the additive term to the sum of the PVE of all genetic terms in two more complex models: one in which a non-additive term was included as a genetic residual following a multivariate normal distribution, and second in which a dominance term following a digenic dominance model was included. As these more complex models did not yield greater PVEs, the simpler model in which only additive effects are included was used for the analyses described below.

A baseline for the reliability of genomic estimated genetic values was obtained by predicting the genetic value for all samples, using the full phenotypic and genotypic datasets available. This form of marker-assisted prediction provides a point of comparison for testing different parameters of the prediction model.

Reliabilities were calculated first as the mean r^2^ returned by StageWise (here denoted as “Stagewise- r^2^”), which is the squared correlation between the true and predicted values assuming the model is correct. This is a model parameter that is proportional to the prediction error variance (PEV) of the BLUP. Second, correlation coefficients between the predicted genotypic values and BLUEs for each genotype calculated on the basis of the observed phenotypes are also reported (here denoted as “Pearson-r^2^”).

### Marker density evaluation

In order to assess the impact of number of SNPs used in the estimation of the variance-covariance “G” matrix by StageWise, the input marker matrix was randomly downsampled to simulate densities ranging from 500 to 45,000 markers, in 100 marker increments. The resulting G matrix was then subtracted from the G matrix calculated using the complete set of markers in order to estimate the ability of smaller marker sets to accurately capture the degree of relatedness between all individuals in each population. Prediction accuracy was also assessed by modeling trait prediction accuracy at different marker densities. At each marker density, twenty random subsets of the marker matrix were produced, with even distribution of markers across the 11 chromosomes, and Pearson-r2 values calculated as described above.

### Population relatedness

The impact of the degree of relatedness between the training and validation population when performing marker-based selection was also assessed, using a leave-one-out cross-validation approach. In the *C. americana* diversity panel, this was done by first sorting the population by the degree of relatedness of each individual in the population to the validation individual, then using a 100-individual subset of the whole population as the training population, which was progressively designed to be less related to the validation individual (i.e., the 1^st^-100^th^ most related individuals, the 2^nd^-101^st^ most related individuals, the 3^rd^-102^nd^ most related individuals, etc.). Once an asymptotically-minimal prediction accuracy was identified, a secondary analysis was performed, by progressively expanding the training population starting from the 20^th^-121^st^ most-related individual, adding in increasing numbers of less-related individuals. This allowed for an assessment of whether training population size could compensate for a lack of training population relatedness.

In the interspecific biparental populations, the known pedigrees of the families provided an alternative method for assessing the impact of relatedness on accuracy. These analyses were performed for each of the three families individually, assessing prediction accuracy by predicting the genotypic value of each member of each of the three full-sib families, using each of the three families as the training population.

## RESULTS

### Morphological in-shell and kernel characteristics

Histograms for six traits, in-shell height and width, kernel height and width, and percent kernel, measured both volumetrically and gravimetrically, are shown in Fig. 2. In general, the interspecific biparental populations, being composed of crosses with *C. avellana* pollen parents from the Oregon State University breeding program, have larger in-shell and kernel dimensions. However, the distribution of percent kernel was observed to be either nearly identical (when measured gravimetrically) or larger in the *C. americana* diversity panel (when measured volumetrically from the digital images).

**Figure 2.**
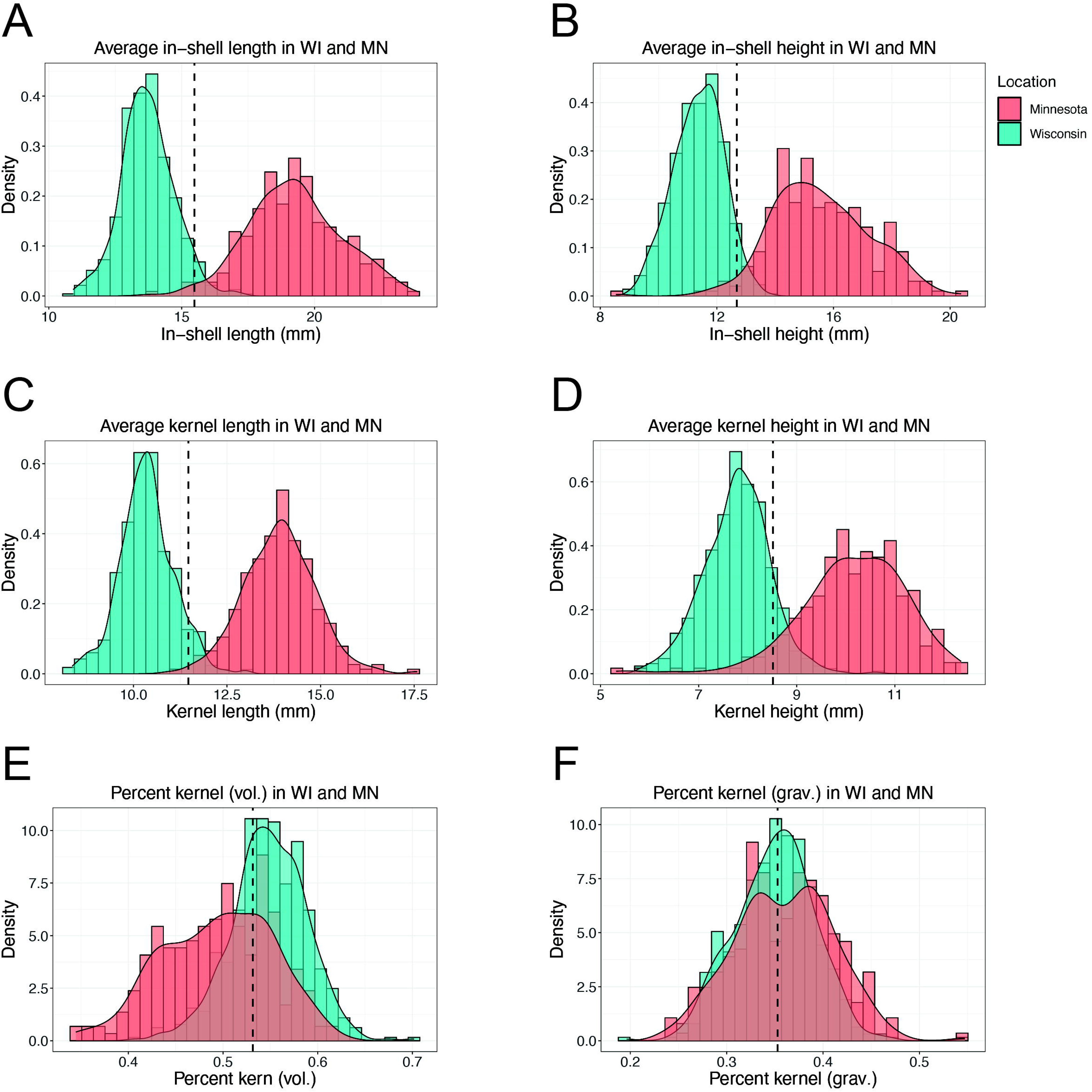
Histograms for six traits measured using digital imagery. Blue distributions correspond to the Wisconsin diversity population of *C. americana*, while red distributions correspond to the interspecific biparental populations in Minnesota. Length and height measurements for in-shell (**A**, **B**) and kernel (**C**, **D**) illustrate the larger size of nuts from the interspecific biparental populations in Minnesota, the paternal parents of which are commercial *C. avellana* cultivars. Slightly greater variance is also observed in the kernel measurements from the interspecific biparental populations. In contrast, higher percent kernel (measured volumetrically) was observed in the *C. americana* diversity population in Wisconsin (**E**), while the two populations had nearly identical distributions for percent kernel when measured by weight (**F**).

Fig. 3 presents PCA biplots for the *C. americana* diversity panel (Fig. 3A), interspecific biparental populations (Fig. 3B) and the combined set of all individuals (Fig. 3C), illustrating the close correlation between in-shell and kernel traits, and their relative independence from both measures of percent kernel.

**Figure 3.**
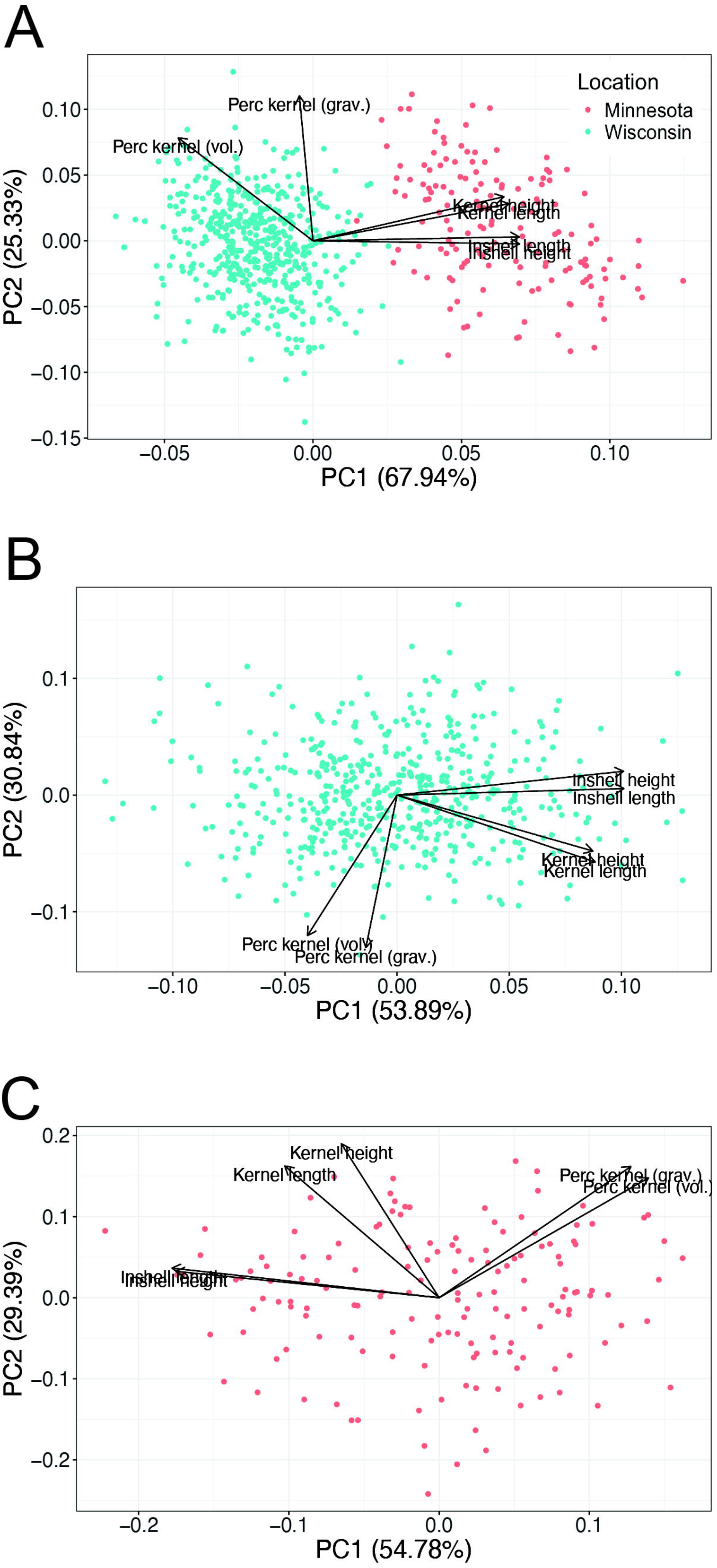
PCA biplots of kernel and nut morphological characteristics, specifically in-shell length, in-shell height, kernel length, kernel height, percentage of kernel by volume, and percentage of kernel by weight. Parallel vectors indicate a positive correlation between the two traits, while vectors oriented 180 degrees from each other indicate negative correlation, and vectors at right angles reflect independence between the relevant traits. In all analyses, kernel height and length, and in-shell height and length, were tightly positively correlated with each other, while volumetric and gravimetric measures of percent kernel were more loosely positively correlated. **A:** In the *C. americana* diversity population in Wisconsin, in-shell and kernel traits were also somewhat positively correlated, and uncorrelated with percent kernel metrics. **B:** In the interspecific biparental populations in Minnesota, the positive correlation between in-shell and kernel measures was substantially reduced, to such a degree that percent kernel metrics approached a negative correlation with in-shell traits. **C:** When both populations were assessed together, correlations appeared much more similar to the *C. americana* diversity panel in Wisconsin (likely due to its larger size), with a slight weakening of the positive correlation between the two measures of percent kernel.

Interestingly, although “flat” nuts (those with a “compression index” – i.e., a width/height ratio – < 0.85) have been reported in some *Corylus americana* germplasm, no such nuts were observed in the present study. The average compression index was 1.09 in the *C. americana* diversity panel, and 1.11 in the interspecific biparental populations, indicating that both groups generally produced round nuts rather than flattened forms.

### Composite interval mapping

The use of multialleic markers in the onemap pipeline produced a high-quality genetic map. Map quality was confirmed by assessing inflation. Given the relatively small size of hazelnut chromosomes (with an average physical size of 29.78 Mb), a given linkage group should not be longer than 100 cM (under normal biological assumptions and a standard mapping function). Inflated map lengths in excess of this upper bound are nevertheless frequently observed by not controlling for genotyping errors, which increase estimates of recombination frequency. Using multi-point regression when converting genetic distances to the final genetic map counteracts this tendency. The linkage groups in our final map are no longer than 89.9 cM, with an average length of 66.3 cM, and a total length of 729.6 cM.

Single QTL scans were performed for each of six phenotypic traits. LOD profiles across the genetic map are shown in Fig 4. Table 1 summarizes the results of the QTL identified via composite interval mapping for each of the six traits. For the two in-shell traits, four QTL were identified (with only one shared in common between both traits), with an average total R^2^ of 52%. For the two kernel-specific measures, three QTL were detected for height, while four were detected for length (with no overlap between the two). In both cases, lower total R^2^ was observed, with an average of 28.6%. The two methods for measuring percent kernel similarly were associated with three (gravimetric) and four (volumetric) unique QTL, with a higher total average R^2^ of 41.1%

**Figure 4.**
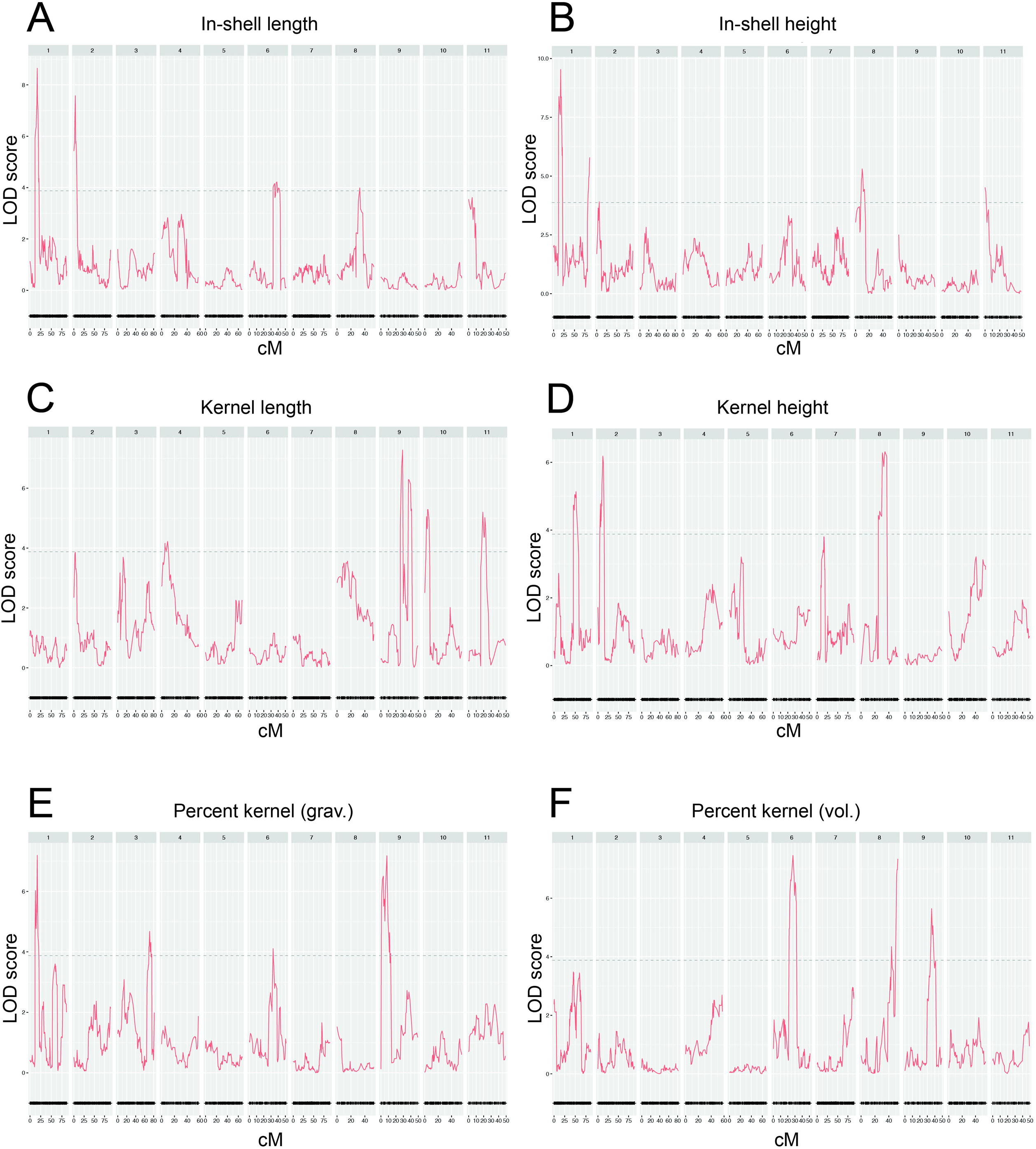
LOD profiles of the composite interval mapping results in the Eric\Jeff interspecific biparental F_1_ population for six morphological nut traits. **A**: in-shell length; **B:** in-shell height; **C:** kernel length; **D:** kernel height; **E:** percent kernel (measured volumetrically); **F:** percent kernel (measure gravimetrically).

**Table 1.**
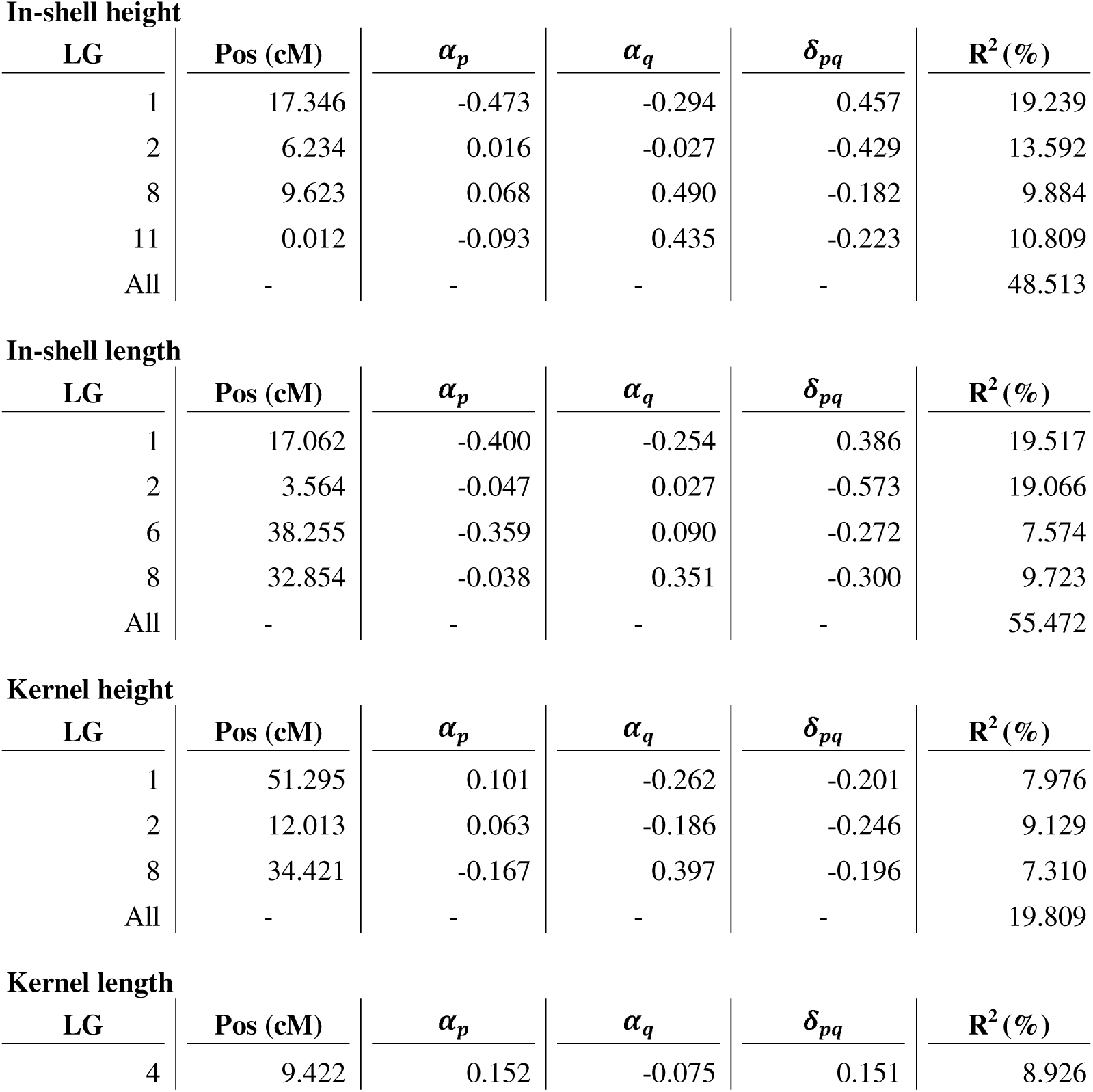
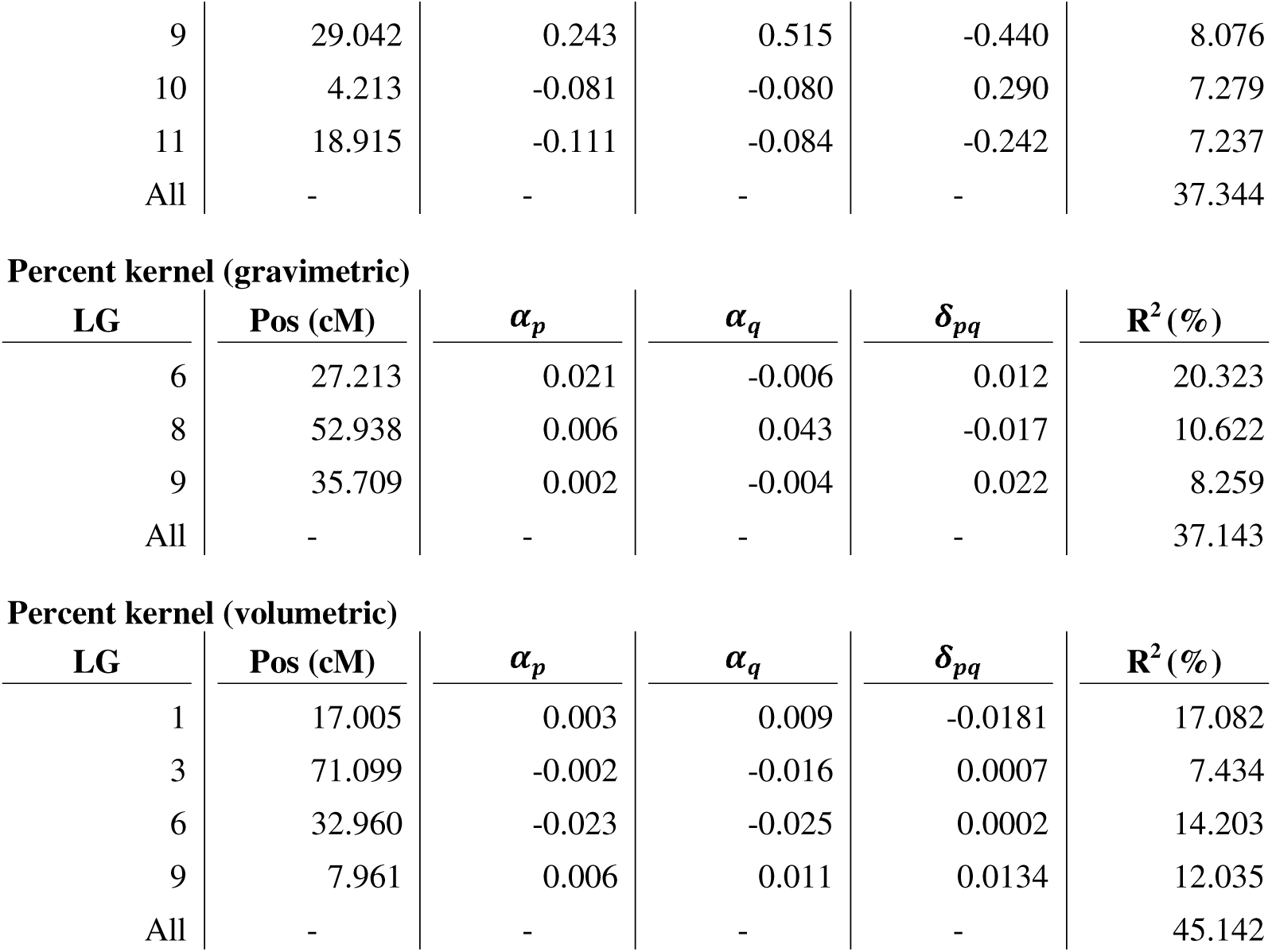
Results of composite interval mapping results in the Eric\Jeff interspecific biparental F_1_ population for six morphological nut traits. For each QTL which exceeded the permutation-test determined LOD threshold, the position is reported in cM along the respective linkage group. In addition, additive effects of both alleles, and dominance effects are reported. R^2^ values for the individual QTL, as well as the complete model are also given.

In-shell height
| LG | Pos (cM) | $\alpha_p$ | $\alpha_q$ | $\delta_{pq}$ | $R^2$ (%) |
| --- | --- | --- | --- | --- | --- |
| 1 | 17.346 | -0.473 | -0.294 | 0.457 | 19.239 |
| 2 | 6.234 | 0.016 | -0.027 | -0.429 | 13.592 |
| 8 | 9.623 | 0.068 | 0.490 | -0.182 | 9.884 |
| 11 | 0.012 | -0.093 | 0.435 | -0.223 | 10.809 |
| All | - | - | - | - | 48.513 |

In-shell length
| LG | Pos (cM) | $\alpha_p$ | $\alpha_q$ | $\delta_{pq}$ | $R^2$ (%) |
| --- | --- | --- | --- | --- | --- |
| 1 | 17.062 | -0.400 | -0.254 | 0.386 | 19.517 |
| 2 | 3.564 | -0.047 | 0.027 | -0.573 | 19.066 |
| 6 | 38.255 | -0.359 | 0.090 | -0.272 | 7.574 |
| 8 | 32.854 | -0.038 | 0.351 | -0.300 | 9.723 |
| All | - | - | - | - | 55.472 |

Kernel height
| LG | Pos (cM) | $\alpha_p$ | $\alpha_q$ | $\delta_{pq}$ | $R^2$ (%) |
| --- | --- | --- | --- | --- | --- |
| 1 | 51.295 | 0.101 | -0.262 | -0.201 | 7.976 |
| 2 | 12.013 | 0.063 | -0.186 | -0.246 | 9.129 |
| 8 | 34.421 | -0.167 | 0.397 | -0.196 | 7.310 |
| All | - | - | - | - | 19.809 |

Kernel length
| LG | Pos (cM) | $\alpha_p$ | $\alpha_q$ | $\delta_{pq}$ | $R^2$ (%) |
| --- | --- | --- | --- | --- | --- |
| 4 | 9.422 | 0.152 | -0.075 | 0.151 | 8.926 |
| 9 | 29.042 | 0.243 | 0.515 | -0.440 | 8.076 |
| 10 | 4.213 | -0.081 | -0.080 | 0.290 | 7.279 |
| 11 | 18.915 | -0.111 | -0.084 | -0.242 | 7.237 |
| All | - | - | - | - | 37.344 |

**Percent kernel (gravimetric)**
| LG | Pos (cM) | $\alpha_p$ | $\alpha_q$ | $\delta_{pq}$ | $R^2$ (%) |
| --- | --- | --- | --- | --- | --- |
| 6 | 27.213 | 0.021 | -0.006 | 0.012 | 20.323 |
| 8 | 52.938 | 0.006 | 0.043 | -0.017 | 10.622 |
| 9 | 35.709 | 0.002 | -0.004 | 0.022 | 8.259 |
| All | - | - | - | - | 37.143 |

**Percent kernel (volumetric)**
| LG | Pos (cM) | $\alpha_p$ | $\alpha_q$ | $\delta_{pq}$ | $R^2$ (%) |
| --- | --- | --- | --- | --- | --- |
| 1 | 17.005 | 0.003 | 0.009 | -0.0181 | 17.082 |
| 3 | 71.099 | -0.002 | -0.016 | 0.0007 | 7.434 |
| 6 | 32.960 | -0.023 | -0.025 | 0.0002 | 14.203 |
| 9 | 7.961 | 0.006 | 0.011 | 0.0134 | 12.035 |
| All | - | - | - | - | 45.142 |

### Genomic prediction accuracy

Two measures of prediction accuracy are presented in Table 2. These are the average StageWise-r^2^ across all individuals in each population, and the Pearson-r^2^, calculated as the correlation between predicted genetic values and phenotypes for all individuals in each population. In the *C. americana* diversity panel, the average StageWise-r^2^ across all traits was 0.56, while the average Pearson-r^2^ was 0.88. In the interspecific biparental populations, these two measures were closer to one another: average StageWise-r^2^ was 0.68, and the average Pearson-r^2^ was 0.67.

**Table 2.** Comparison of prediction accuracies for six nut and kernel traits in the *C. americana* (Wisconsin) and interspecific hybrid (Minnesota) populations. StageWise-r² values represent model-derived prediction reliability based on genetic variance components, while Pearson-r² values reflect the squared correlation between predicted and observed phenotypic BLUEs. Together, these metrics summarize the performance of genomic prediction models across traits and populations.

| Trait | Wisconsin –<br>StageWise-$r^2$ | Wisconsin –<br>Pearson-$r^2$ | Minnesota –<br>StageWise-$r^2$ | Minnesota –<br>Pearson-$r^2$ |
| --- | --- | --- | --- | --- |
| Height (in-shell) | 0.55 | 0.90 | 0.88 | 0.87 |
| Length (in-shell) | 0.51 | 0.85 | 0.84 | 0.75 |
| Height (kernel) | 0.62 | 0.93 | 0.44 | 0.51 |
| Length (kernel) | 0.53 | 0.83 | 0.48 | 0.42 |
| % kernel (grav.) | 0.61 | 0.92 | 0.74 | 0.75 |
| % kern (vol.) | 0.53 | 0.87 | 0.71 | 0.73 |

### Effect of marker density on prediction accuracy

The results of the marker density analysis are shown in Fig. 5, where the y-axis represents the absolute value of the sum of the matrix resulting from the subtraction of the downsampling- derived G matrix. In both populations, an asymptotic minimum in the difference between the matrices is reached between 5,000 and 10,000 markers.

**Figure 5.**
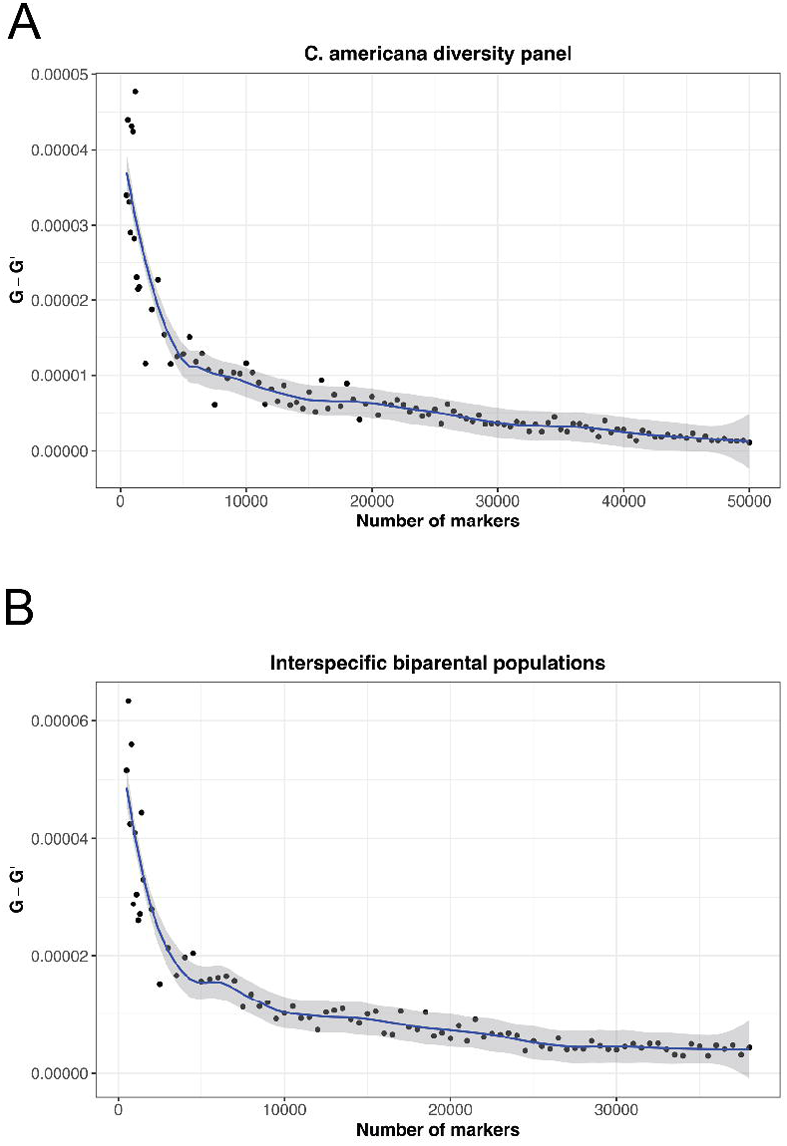
Assessment of the impact of marker density on pairwise estimates relatedness within both the *C. americana* diversity panel, and the interspecific biparental populations. The **G** matrix was calculated with the complete marker set, and then for each number of markers along the *x*-axis, a **G** matrix was calculated again using StageWise. This latter matrix was then subtracted from the original G matrix to give a scalar difference, representing the deviation in total estimates of relatedness at varying levels of marker density. **A:** results for the *C. americana* diversity panel in Wisconsin; **B:** results for the interspecific biparental populations in Minnesota.

A similar pattern was observed for average StageWise-r^2^ values, where there was a rapid increase in mean StageWise-r^2^ as marker density increased, until an asymptotic maximum StageWise-r^2^ was attained between 5,000 and 10,000 markers (Supp. Fig. 2), further suggesting that a marker density in this range will most efficiently estimate relatedness between individuals. When all individuals were used in the calculation of BLUPs, the difference was relatively minor, representing an increase of only ∼0.04 in the case of the *C. americana* diversity population. In the interspecific biparental populations, the difference is almost entirely negligible: while the exponential relationship is evident, the difference between the minimum and maximum reliability is ∼0.01.

### Impact of relatedness on prediction accuracy

Results from the comparison of variably-related training populations are shown in Fig 6. Prediction accuracy quickly drops as the most closely related individuals are removed from the training population, leading to asymptotically-minimal accuracies when the twenty most-closely related individuals are removed. This reduction in accuracy represented a nearly 55% reduction in relatedness (measured by the average of the coefficients of the corresponding elements of the G matrix). However, even at maximal relatedness, prediction accuracy in the *C. americana* diversity population was relatively low, with maximum Pearson-r^2^ values of ∼0.45, and an average of 0.3.

**Figure 6.**
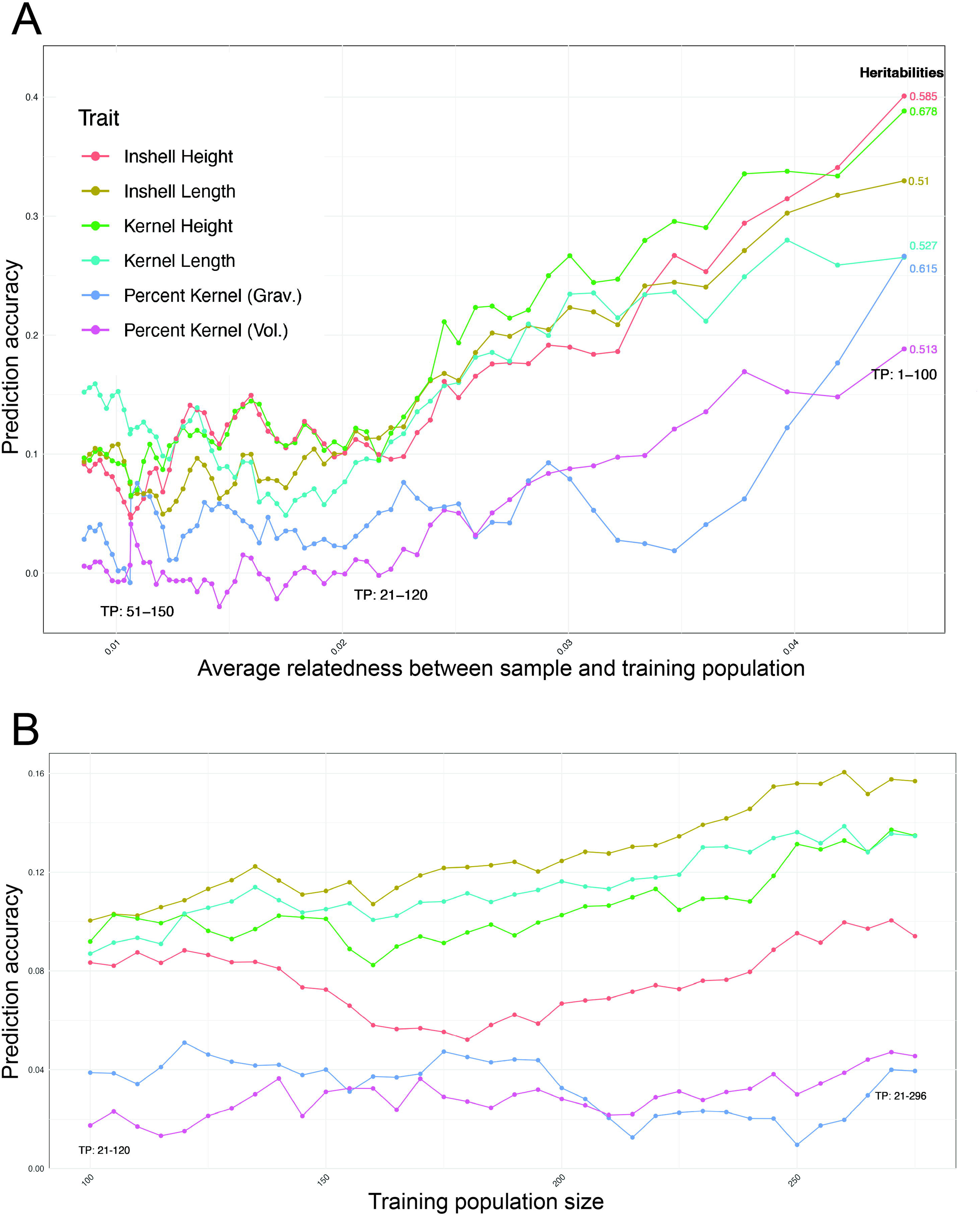
The impact of training population design on prediction accuracy in the *C. americana* diversity panel in Wisconsin. **A:** The impact of decreasing the relatedness of the training population to the predicted sample. Declining prediction accuracy is observed as the most related individuals are removed from the training population, until an asymptotically minimum prediction accuracy is reached when the twenty most-closely related individuals have been removed. **B:** The impact of increasing population size on prediction accuracy, when these twenty most closely related individuals have been removed. Population size was increased from 100 to 275 individuals, with marginal increases in prediction accuracy observed for four traits (in-shell and kernel height, in-shell and kernel length), while prediction accuracy for both measures of percent kernel were unaffected.

The ability to compensate for reduced prediction accuracy by increasing the size of the training population was subsequently assessed. The results of this analysis are shown in Fig. 6. For the two measures of percent kernel (which also exhibited the lowest prediction accuracy in general), this increase in population size had no impact on prediction accuracy, while for in-shell height, the reduction in relatedness associated with increasing population size led to a reduction in prediction accuracy. Finally, for kernel height, kernel length, and in-shell length, prediction accuracy slightly increased, however all of these changes in prediction accuracy represented less than 0.25% in total variation. This indicates that if the 20 most-closely related individuals are absent from the training population design, this cannot be substantially offset by simply including more weakly-related individuals. Importantly, the average degree of relatedness in the *C. americana* panel is relatively low, and thus this observation is not necessarily generalizable to all other training population designs.

In the interspecific biparental populations, similar results were observed when reducing the relatedness of the training population, and keeping the total size of the population constant at 100 individuals Supp. Fig. 3. However, there are limitations to this approach for these populations. Because of the structured nature of the full-sib families, correlations between the genotypic values and phenotypic data across all individuals primarily reflects this structure. These results are shown in Table 3. Interestingly, marker-based prediction led to low (near-zero or negative) Pearson-r^2^ values, when the training population represented the same interspecific biparental cross from which the sample being validated was drawn. For each of the families, however, there was one other biparental population which led to positive prediction accuracies, although these maximum Pearson-r^2^ values were significantly lower than the maximum values observed in the *C. americana* diversity panel.

**Table 3.** Impact of relatedness on prediction accuracy in the interspecific biparental populations in Minnesota. For each of the three F_1_ families, leave-one-out validation was performed using the entirety of each of the three families, with average Pearson-r^2^ values being calculated. The prediction population which generated the highest prediction accuracy is bolded.

| Sample population | Prediction Population | Prediction accuracy |
| --- | --- | --- |
| Gibs\York | Eric\Jeff | -0.128 |
| **Gibs\York** | **Gibs\OSU** | **0.281** |
| Gibs\York | Gibs\York | -0.0545 |
| Eric\Jeff | Eric\Jeff | -0.723 |
| **Eric\Jeff** | **Gibs\OSU** | **0.404** |
| Eric\Jeff | Gibs\York | -0.0219 |
| **Gibs\OSU** | **Eric\Jeff** | **0.245** |
| Gibs\OSU | Gibs\OSU | -0.638 |
| Gibs\OSU | Gibs\York | 0.220 |

## DISCUSSION

### Quantitative variation and few large-effect QTL for morphological nut traits

The composite interval mapping results reported above demonstrate clearly that the traits measured here related to physical characteristics of in-shell nuts and kernels are highly quantitative and likely polygenic. We have observed moderate to high heritability for these traits, yet the majority of phenotypic variance remains unexplained by the limited number of detected QTL. This distinction emphasizes that while a few loci of measurable effect can be identified through CIM, much of the underlying genetic variance likely arises from numerous small-effect loci. On average, the identified QTL individually explained only 11% of the phenotypic variation for a given trait. While for certain traits the total explained variance was much higher, the average total R^2^ for all QTL associated with a trait was only 40.5%, with the unexplained variance ranging from 44.5%-80.2%. A low total R^2^ was particularly apparent for kernel traits, which was also reflected in the prediction accuracy for these traits when evaluating genomic prediction models using the interspecific biparental populations in Minnesota.

To our knowledge, this is the first study to ever map kernel and nut quality traits in an interspecific *C. americana* x *C. avellana* family. Several studies have performed association analysis in *C. avellana* populations of similar size. Baytar et al. (2024) used similar GBS marker data, and evaluated both kernel and nut size, as well as percent kernel. However, they found only a single QTL for percent kernel. Another GWAS utilized a low density SSR map and only found one QTL for kernel weight and none for percent kernel (Ozturk et al., 2017). Most comparable to these results was a linkage mapping study performed using GBS markers and a relatively large F1 population derived from Italian *C. avellana* cultivars (Torello Marinoni et al., 2018). This analysis included phenotypic data on both nut and kernel size, identifying one QTL for kernel size (on LG 5) and four for nut size (on LGs 1, 2, 4, and 6). However, none of these QTL explained more than 10% of the total phenotypic variance.

In light of these results, the CIM analyses reported here represent a significant advance in our understanding of the genetic control of these physical nut characteristics, which is likely attributable to the significant variation observed in these interspecific crosses, the high resolution of the genetic map used, and the precision and scale of the phenotyping method utilized.

Nevertheless, it appears likely that this study was underpowered to detect the numerous remaining QTL associated with these traits. This is likely not due to a limitation in the number of markers used, since the resolution of the map was greater than the number of expected recombination events present in the population. Instead, it is likely that the relatively small size of the mapping population used in this study is primarily responsible for reduced statistical power. Building larger F_1_ populations would therefore be useful in attempting to more accurately assess the true genetic architecture of these traits, however, the size of F_1_ progeny families are limited by *Corylus* biology. Multiparent QTL mapping would therefore be a valuable approach (e.g., via a diallel mating design), as this would enable more precise estimation of dominance effects. Nevertheless, the present results point strongly towards many QTL influencing these traits, and the general absence of very large effect QTL. As will be discussed below, this motivates relying upon genomic prediction models as the preferred method for utilizing marker information in applied breeding programs.

### Discrepancies between the *C. americana* diversity panel and interspecific biparental populations

While the correlation coefficient-based estimates of prediction accuracy in the interspecific biparental populations were nearly identical (but slightly less than) the mean Pearson-r^2^ values, correlation coefficients in the *C. americana* diversity population were dramatically higher than the mean Pearson-r^2^ values. This difference is so great that while mean Pearson-r^2^ values were higher in the interspecific biparental populations than in the *C. americana* diversity population (higher for every trait other the kernel-specific measurements), correlation coefficients were much higher in the *C. americana* diversity population than in the interspecific biparental populations. Possible interpretations of these discrepancies, and how they should inform the interpretation of these two metrics of reliability, are provided below. It should be noted at the outset that because best linear predictors are designed to maximize the correlation between the true and predicted genotypic values (and the covariance of the true and predicted values is equal to the variance of the true values), reliability of the model is equivalent to broad sense heritability as computed in a completely randomized design (Endelman, 2023). Correlation coefficients derived from a leave-one-out cross-validation approach can introduce bias by assuming the observed phenotypic data represents the “true” value of an individual (Zhou et al., 2017). However, in general, it is clear that marker-assisted selection generated strikingly high reliabilities, particularly for populations composed of unreplicated seedling genotypes. Except for the kernel-specific traits in the interspecific biparental populations, both measures of reliability were higher than 0.5. While to our knowledge this study represents the first reported evaluation of genomic selection in any *Corylus* species, these results are nevertheless comparable to assessments of prediction accuracy for morphological characteristics in other tree nut crops. For example, Nishio et al. (2018) reported prediction accuracies of 0.4 for the weight of chestnuts using a BayesB model.

There are numerous differences between the *C. americana* diversity population and the interspecific biparental populations in Minnesota which likely underlie the differences in trait heritabilities observed for these two locations. While spatial variation within the field locations was not apparently greater in either the *C. americana* diversity population or the interspecific biparental populations (Supp. Fig. 4), the fact that both trials were composed of unreplicated seedling genotypes certainly lowered heritabilities, and unavoidably increased the relative size of environmental variance components, potentially to differing degrees.

In addition, the populations themselves were obviously markedly different. While the interspecific biparental populations were substantially smaller, they also exhibited much higher degrees of average relatedness both within and across biparental populations. The *C. americana* diversity population location was composed of wild-collected accessions with no known shared pedigree. The interspecific biparental populations, on the other hand, were composed of full-sib crosses in which maternal parents were both drawn from the Minnesota breeding program, and paternal parents were selected from the Oregon breeding program. As a result, in most training populations designed to test prediction accuracy in the interspecific biparental populations, the average coefficients of the respective G matrix were roughly an order of magnitude higher than in the *C. americana* diversity panel.

### Excessive shrinkage within the interspecific biparental populations

The cross-validation results analyzing the impact of relatedness on prediction accuracy in the interspecific biparental populations are striking, and are consistent across traits. It is not immediately clear why prediction accuracies were consistently negative when the training population was composed of the same full-sib family as the individual for which the prediction was being made. This is particularly surprising, given the relatively high accuracies (which for certain traits, exceeded the asymptotic maximum prediction accuracy seen in the *C. americana* diversity population) when the prediction population selected is **not** the full sib family from which the given sample is drawn. Two important features of this analysis are likely impacting these results. First, there is substantial shrinkage, much greater than that seen in the *C. americana* diversity population, apparent across traits when performing these cross-validation analyses (Supp. Fig. 5). It has been previously reported that shrinkage-based estimators can lower prediction accuracy for unphenotyped individuals when predictions are made in small populations with unreplicated genotypes (Endelman and Jannink, 2012; Zhao et al., 2013).

Second, and relatedly, the population sizes for these comparisons across families is small, ranging from 75-81 individuals being used in the given prediction populations. This is much smaller than is typically considered in studies assessing factors influencing genomic prediction accuracy, where significant drop-offs in accuracy are frequently observed when population size drops below 100 individuals (Tayeh et al., 2015; Zhang et al., 2017; Sverrisdóttir et al., 2018; Edwards et al., 2019). These were also smaller training populations than those used to analyze the impact of relatedness in the *C. americana* diversity population. Generating larger biparental families is biologically challenging in hazelnut, so breeding programs should adopt a strategy of ensuring biparental families are interconnected through shared parentage, such that higher effective population sizes can be achieved in training populations.

### Mean r^2^ values vs. correlation coefficients as measures of reliability

The average of the r^2^ values returned by the StageWise package and the Pearson correlation coefficients between the genotypic values estimated by StageWise and the BLUEs represent two intuitive methods for estimating reliability. The high values for reliability reported here (>0.5 in nearly all cases) suggest relatively large broad-sense heritabilities i.e., large additive genetic variance components, for morphological nut traits. This suggests that applying genomic prediction methods within selection programs is realistic and tenable. There are, however, several caveats to be aware of in evaluating these measures of reliability.

Any correlation-based measure of reliability will be inflated to a degree that is proportional to the correlation between population structure and the traits of interest (Werner et al., 2020). While predicated on the assumption that the model is “correct”, and thus potential underestimates of “true” reliability, the mean r^2^ of the BLUPs returned by StageWise should in principle be less sensitive to this artificial inflation, as they are individual measures of correlation across replicates of each specific genotype (Endelman, 2023). At the same time, these correlation coefficients will only ever be accurate measures of true reliability to the degree that the BLUEs are taken as a form of “groundtruthed” data (Rincent et al., 2012). In the case of unreplicated seedling genotypes, where environmental variance components cannot be explicitly included in the model, this assumption is clearly not entirely justified (Waldmann, 2019), leading to an artificial reduction in Pearson-r^2^ values from what might be observed in, e.g., an analysis of an RCBD.

### Application of genomic prediction within hazelnut breeding programs

There are numerous practical considerations that will impact the use of genomic prediction in the various stages of a specific hazelnut breeding program. In particular, the per sample cost of genotyping will be a determining factor influencing the number of plants for which predictions can be made. In addition, important traits may express themselves relatively quickly (e.g., via disease screens which can be performed in a greenhouse environment), or take over a decade before a phenotype is available (e.g., mature plant architecture, or multi-year cumulative yield). This will significantly impact the benefit of genomic-based selection, relative to phenotypic selection.

In general, however, several conclusions are evident from the data presented here. First, prediction accuracies that are obtained when phenotypic data is not masked for individuals in the training population are clearly high enough to warrant including genetic marker information in the selection of parents. The set of parental candidates is typically relatively small, and breeders frequently have limited access to replicated trial data for all individuals with which crosses might realistically be made. At the same time, some phenotypic data will be available before a cross can physiologically be performed, and thus the inclusion of genotypic information would in this context be used to perform genomic-assisted selection (Jannink et al., 2010). In addition, in mature breeding programs, close relatives of potential parents may have substantial phenotypic information available, which can be used to improve the training population. The improvements in trait prediction offered by such an approach will generally be warranted, given historically- consistent declines in the cost of sequencing.

Second, in many applied contexts, genomic prediction is not simply used in a marker-assisted fashion, but also to perform marker-based prediction, where only genotypic data is available for some individuals in the population. In these cases, a so-called “training population” is phenotyped and genotyped, allowing genotypic values to be predicted for un-phenotyped individuals. The use of purely genomic-based prediction at the seedling stage, performed to determine which subset of progeny families to advance into field trials, will necessarily be more limited. Phenotypic data will be largely absent for such individuals. While there may be cases in which phenotypic data exists for half-sib or full-sib families created from crosses using the same parents, any training population will be to some degree removed from the individuals for which predictions are being made. In addition, while the size of any specific progeny family will be limited, a breeding program as a whole could easily generate thousands of seedlings each year, genotyping all of which would likely represent a prohibitive expense. Instead, a prudent balancing act should be made between resources dedicated to genotyping, and field trials used to grow out new progeny each year, such that the number of progeny generated each year can in fact be genotyped. Even in the absence of phenotypic data, and thus reduced prediction accuracies, genotyping at the seedling stage will improve genetic gain, and allow for a more informed preservation of genetic variance within the breeding program (Endelman, 2025). While fixed-site marker platforms such as KASP assays offer a lower-cost alternative for validating specific QTL (Semagn et al., 2014), they are generally insufficient for maintaining prediction accuracy in complex, polygenic traits where genome-wide coverage is required. Emerging targeted amplicon sequencing approaches may provide a cost-effective compromise, offering sufficient marker density for genomic prediction while reducing per-sample genotyping costs relative to GBS or array-based platforms (Dobosy et al., 2011; Kilian et al., 2012).

In sum, these results provide a promising basis for further developing genomic selection models for use in hazelnut breeding programs. This study represents the first such model developed in the *Corylus* genus, and despite the limited degree of replication, the prediction accuracies obtained for nut quality traits suggest this method can substantially improve the rates of genetic gain for polygenic traits. We hope future studies will expand upon this work, validating models with more robust experimental designs to better minimize environmental variances, and validate predication accuracies for a greater diversity of traits.

## ACKNOWLEDGEMENTS

The Dawson Lab assisted greatly in the collection of phenotypic and genotypic data, specifically, Marissa Nix, Peyton Higgins, Martha Barta, Ava Gorius, Ava Glaser, Tressa Peskar, Malachi Persche, Matt Mirkes, Shea Tillotson, Raeann Rich, Maya Giordano, Calliana Wickus, Thomas Hickey, and Brent Johnson. We are thankful to the support of Chuck and Gerta Zinda, for allowing us access to the *C. americana* diversity panel in Wisconsin, as well as Mark Hamann, Lois Braun, and Les Everett, for providing the interspecific biparental families in Minnesota.

## SUPPLEMENTAL MATERIAL

**Supplementary Figure 1.** Comparison between hand-measured and image-derived phenotypic values for nut and kernel traits. Each point represents a mean value per individual bush, with dashed lines indicating 1:1 correspondence. Digital measurements were highly correlated with manual caliper measurements across all traits, confirming the accuracy of the image-based phenotyping platform.

**Supplementary Figure 2.** The impact of marker dilution is illustrated in terms of its impact on prediction accuracy, as measured by Pearson-r^2^ values. **A:** results from the *C. americana* diversity panel; **B:** results from the interspecific biparental populations. As in Fig. 5, asymptotic prediction accuracy is reached between 5,000 and 10,000 markers.

**Supplementary Figure 3.** Progressive dilution of the training population is shown for the interspecific biparental populations in Minnesota. The analysis is similar to what is shown in Fig. 6, however in this case, prediction accuracy is largely a function of correctly predicting F_1_ family membership, and not the genetic value of individual samples, due to the significant degree of population structure present when all three biparental families are combined.

**Supplementary Figure 4.** Spatial variation across the field sites for **A:** kernel length in the Wisconsin *C. americana* diversity panel; **B:** kernel height in the Wisconsin *C. americana* diversity panel; **C:** kernel length in the Minnesota interspecific biparental populations; **D:** kernel height in the Minnesota interspecific biparental populations. No clear structured spatial variation is observed within sites, or any degree of relatively greater or lesser variation between sites.

**Supplementary Figure 5.** Violin plots comparing observed phenotypic values with the genetic values predicted using BLUPs. **A:** Shrinkage is observed in the *C. americana* diversity panel, and this effect increases as the training population becomes less related to the sample. **B:** Much more extreme shrinkage is observed in the biparental populations across all traits, despite the similar training population size.

## DATA AVAILABILITY

Digital imagery (both raw and binary masks) which were used for phenotyping, genetic map using multialleic makers, ordered linkage groups for the “Eric\Jeff” F_1_ population, input marker and phenotype files for use in fullsibQTL, and input marker and phenotype files for use in StageWise, are available via DataDryad [temporary link for peer review: http://datadryad.org/share/DdbiZHYdCypYeb2d7gCfkZC8edWZ4jpWjfmRZcyitpw].

## Abbreviations

EFB: Eastern Filbert Blight, *Anisogramma anomala*
UMHDI: Upper Midwest Hazelnut Development Initiative
Eric\Jeff: Eric4-21 x Jefferson
Gibs\OSU: Gibs5-15 x OSU-919-031
Gibs\York: Gibs5-15 x York
LD: linkage disequilibrium
RSS: residual sums of squares
BLUPs: best linear unbiased predictors
BLUEs: best linear unbiased estimators
PVE: percent variance explained
Stagewise- r2: mean r2 returned by StageWise
Pearson-r2: correlation coefficients between the predicted genotypic values and BLUEs for each genotype calculated on the basis of the observed phenotypes
QTL: quantitative trait loci
LG: linkage group
SNP: single nucleotide polymorphism

