## Supplementary figures and images for "Composite interval mapping and genomic prediction of nut quality traits in American and American-European interspecific hybrid hazelnutss"

### Supplementary Figure 1

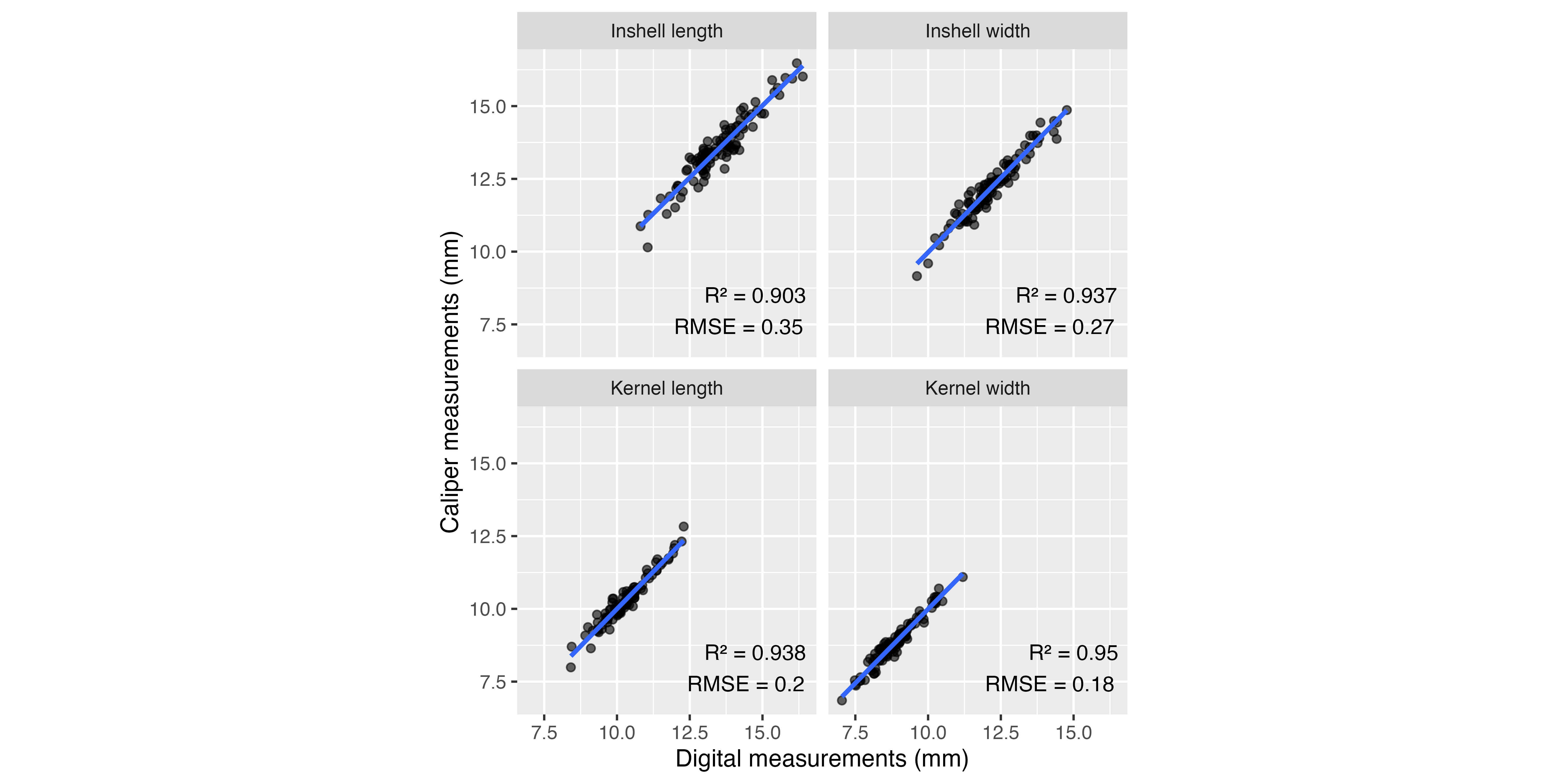

### Supplementary Figure 2

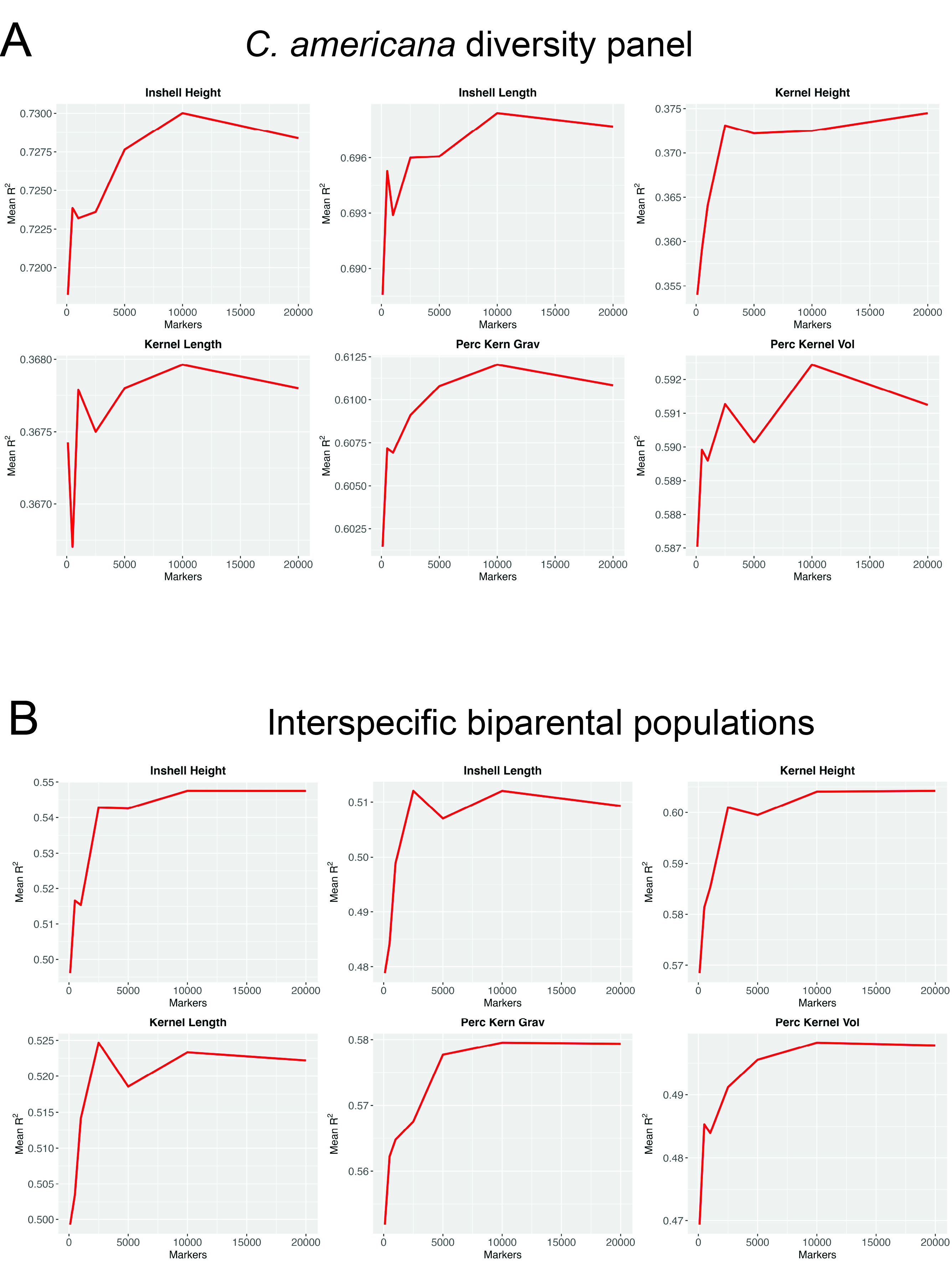

### Supplementary Figure 3

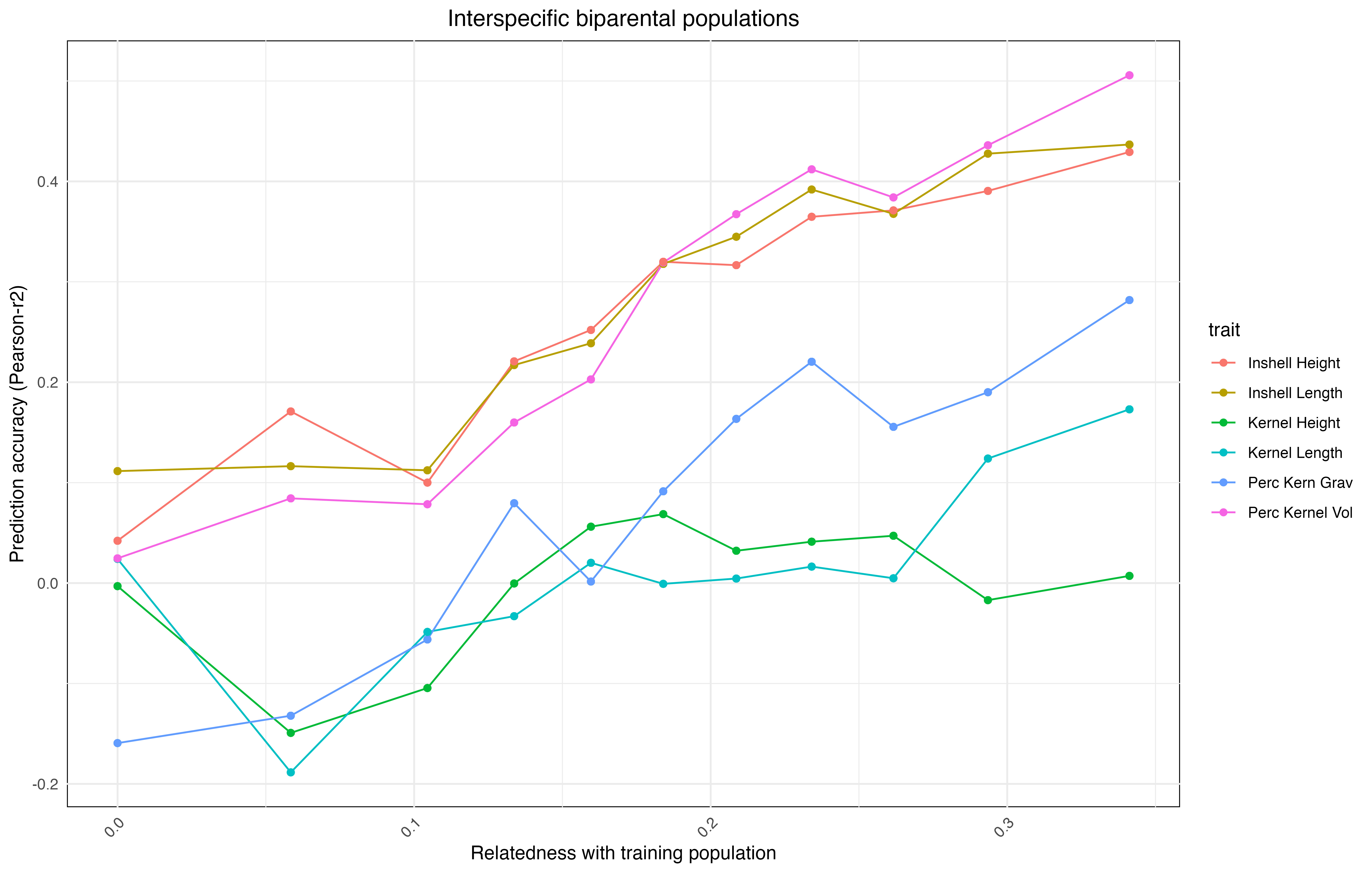

### Supplementary Figure 4

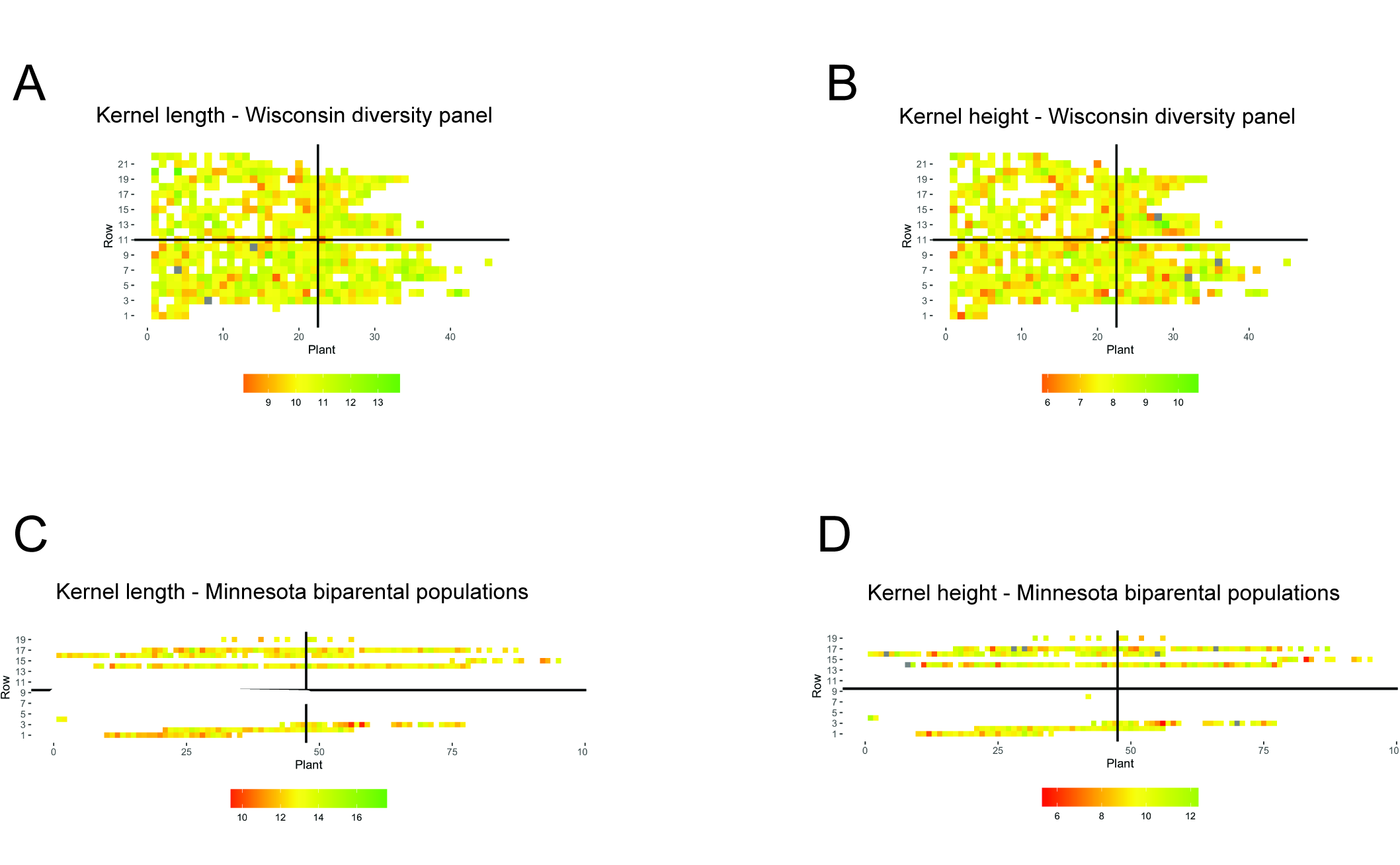

### Supplementary Figure 5

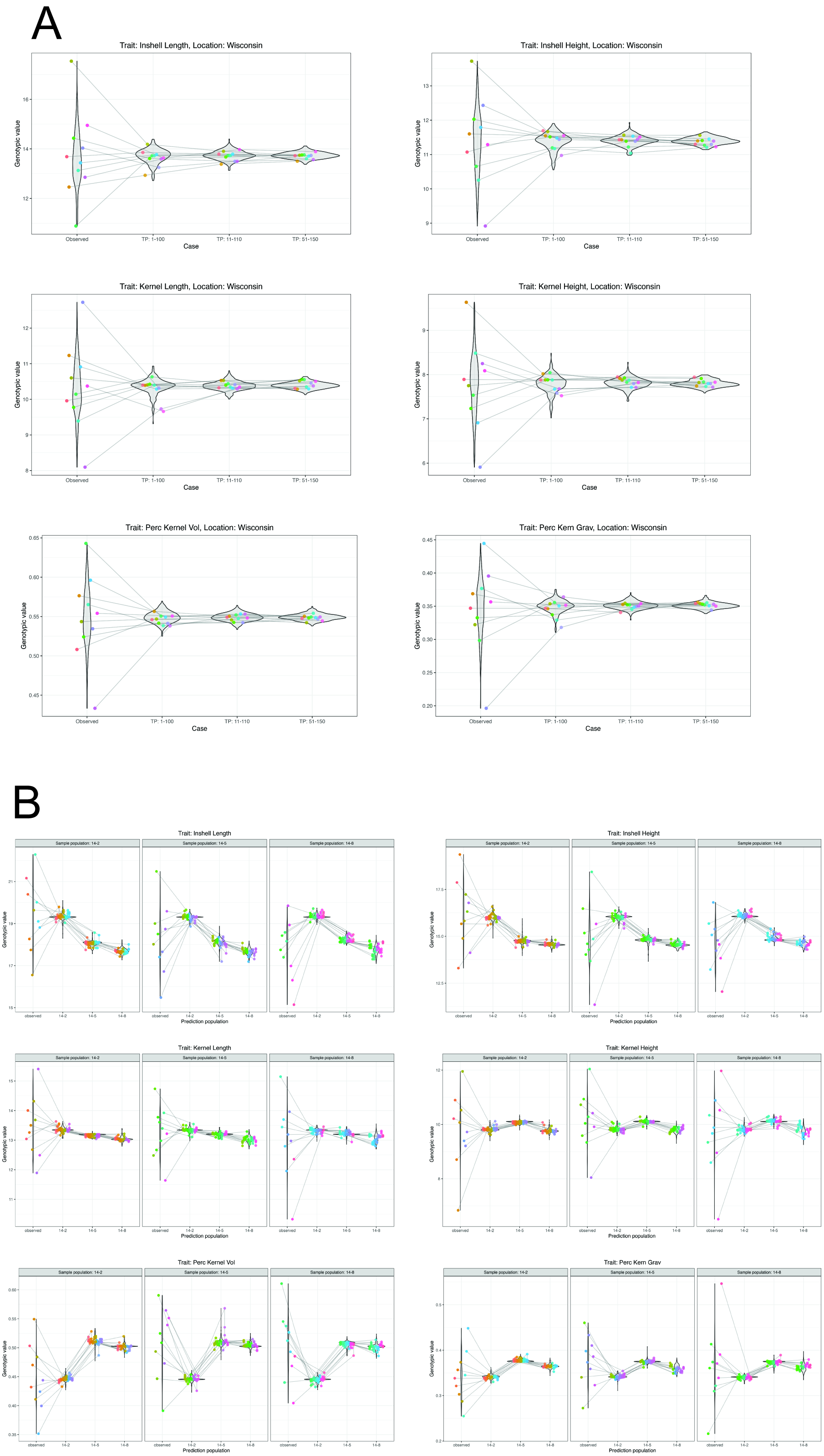
